# Genome-wide post-transcriptional dysregulation by microRNAs in human asthma as revealed by Frac-seq

**DOI:** 10.1101/234500

**Authors:** Rocio T. Martinez-Nunez, Hitasha Rupani, Manuela Platé, Mahesan Niranjan, Rachel C. Chambers, Peter H. Howarth, Tilman Sanchez-Elsner

## Abstract

MicroRNAs are small non-coding RNAs that inhibit gene expression post-transcriptionally, implicated in virtually all biological processes. Although the effect of individual microRNAs is generally studied, the genome-wide role of multiple microRNAs is less investigated. We assessed paired genome-wide expression of microRNAs with total (cytoplasmic) and translational (polyribosome-bound) mRNA levels employing Frac-seq in human primary bronchoepithelium from healthy controls and severe asthmatics. Severe asthma is a chronic inflammatory disease of the airways characterized by poor response to therapy. We found genes (=all isoforms of a gene) and mRNA isoforms differentially expressed in asthma, with novel inflammatory and structural mechanisms disclosed solely by polyribosome-bound mRNAs. Gene expression (=all isoforms of a gene) and mRNA expression analysis revealed different molecular candidates and biological pathways, with differentially expressed polyribosome-bound and total mRNAs also showing little overlap. We reveal a hub of six dysregulated microRNAs accounting for ∼90% of all microRNA targeting, displaying preference for polyribosome-bound mRNAs. Transfection of this hub in healthy cells mimicked asthma characteristics. Our work demonstrates extensive post-transcriptional gene dysregulation in asthma, where microRNAs play a central role, illustrating the feasibility and importance of assessing post-transcriptional gene expression when investigating human disease.

## INTRODUCTION

MicroRNAs are small regulatory molecules (∼22 nucleotides long) that inhibit gene expression by pairing mainly to the 3’ UTRs (UnTranslated Region) of their target mRNAs (1). MicroRNA effects include mRNA destabilization and inhibition of translation, with a body of literature supporting both as main effector mechanisms (2-5). The biological relevance of microRNAs expands to most cellular processes, as thousands of mRNAs contain microRNA responsive elements (MREs). Consequently, microRNA dysregulation has been demonstrated to underlie disease pathophysiological mechanisms, making microRNAs novel therapeutic targets (6). Although the effects of individual microRNAs in disease have been explored extensively, there are fewer reports on the role of microRNAs acting as networks in this setting. We have previously shown that microRNAs dysregulated in asthmatic bronchial epithelial cells may have different effects-even opposite-when modulating their levels individually *vs* simultaneously (7). Our work and others (8) highlight the need for integrative genome-wide approaches to understand the role and importance of microRNAs in cellular and pathological processes.

Asthma is a common chronic inflammatory disease of the airways affecting ∼350 million people worldwide and with a spectrum of severity. Severe asthma (SA) is characterized by the need for or the failure to respond to high dose glucocorticoids in conjunction with other additional controller therapies (9). The underlying mechanisms of severe asthma remain incompletely understood and it therefore represents a major unmet clinical need, accounting for the majority of the healthcare budget dedicated to asthma. Since severe asthma patients remain uncontrolled it is probable that additional mechanisms or steroid-unresponsive processes contribute to the disease persistence. As the airway epithelium orchestrates both inflammatory and remodeling processes relevant to severe asthma (10,11), we have focused on investigating alterations in this cell population. Moreover, we have centered on investigating post-transcriptional control of gene expression in asthma given that post-transcriptional control is considered key in the regulation of inflammation (12) and requires further understanding in many diseases including asthma.

Popular approaches to studying complex diseases using high throughput methodologies focus on the transcriptome, measured with arrays and sequencing technologies. However, it is appreciated that gene transcription and gene translation are not synonymous. From transcription to translation into protein, mRNA undergoes multiple processes including splicing, stabilization, targeting by microRNAs and decay (13) which affect mRNA loading into polyribosomes and subsequent translation. It is well acknowledged that the transcriptome shows weak correlation with the corresponding protein levels, as has been noted in early work (14). More recent works (15-17) show that the weak correlation can be improved upon by a machine learning approach integrating transcript levels, transcript stability, polyribosome binding and other sequence-based proxies of translation rates. Thus, the disparity between mRNA and protein levels in a variety of systems reflects the relevance of post-transcriptional mRNA regulation, demonstrating that cytoplasmic mRNA expression inadequately reflects actual translation into protein (13,18). This disparity is even more pronounced in mammalian cells because of mRNA splicing. Alternative splicing generates several mRNAs from one single gene, with virtually all genes undergoing splicing (19). Alternatively spliced mRNA isoforms show preferential binding to polyribosomes (20) and heavily influence protein levels (21). Together these observations highlight the need for consideration of splicing and translation when performing genome-wide mRNA expression measurements.

Given that microRNAs may affect mRNA levels and/or their translation into protein, as well as their importance in disease, we sought to determine the genome-wide relationship between microRNAs and their mRNA targets in human asthma in different sub-cellular compartments. To this end, we performed microRNA profiling using small RNA-sequencing and integrated it with Frac-seq (20) in bronchial epithelial cells (BECs) isolated from human clinical samples from healthy volunteers and well-characterized severe asthmatic patients. Frac-seq combines subcellular <u>frac</u>tionation and RNA <u>seq</u>uencing, measuring mRNA levels in both cytoplasm (all mRNAs) and polyribosome bound (mRNAs undergoing translation) fractions (22), facilitating the study of post-transcriptional mRNA regulation on a genome-wide scale.

Our work presents for the first time evidence that genome-wide control of mRNA splicing and translation is at the centre of a human disease and asthma pathophysiology. Our results show that microRNAs preferentially target polyribosome-associated mRNAs in asthma, adding valuable knowledge to the long-standing debate about microRNA effects on their targets (2,5,23,24). More strikingly, amongst the microRNAs detected as differentially expressed between health and asthma, our results show that ∼50% of the changes in mRNA binding to polyribosomes is modulated by a small hub of only six microRNAs. These six microRNAs account for ∼90% of all cellular microRNA targeting and recapitulate disease characteristics when modulated in healthy cells. Our work highlights the relevance of studying microRNAs in their molecular and cellular context and demonstrates the feasibility and importance of studying post-transcriptional gene regulation in human disease, opening a novel path in the understanding of the asthmatic process and potentially other pathologies.

## RESULTS

### Severe asthma patients present genome-wide differences in microRNA levels compared to healthy donors as determined by small RNA-seq

To determine microRNA expression we performed small RNA-sequencing on BECs (HC=5, SA=8, Supplementary Table S1). Principal Component Analysis (PCA) (P < 0.05, 0.66 > SA/HC ratio > 1.5, Supplementary Table S2) identified that genome-wide microRNA expression is different between HC and SA patients. Unsupervised hierarchical clustering of 21 differentially expressed microRNAs separated the samples between health and disease, with the exception of SA3 (Figure 1B). We validated these findings with microRNA RT-qPCRs on an expanded cohort (n=9 HC and n=11 SA, Figure 1C, most differentially expressed microRNAs). MicroRNAs −19b-3p, −20a-5p, 135b-5p, −574-3p and 625-3p were validated, while only miR-127-3p showed no significant differences between SA and HC amongst the candidates tested.

**Figure 1.**
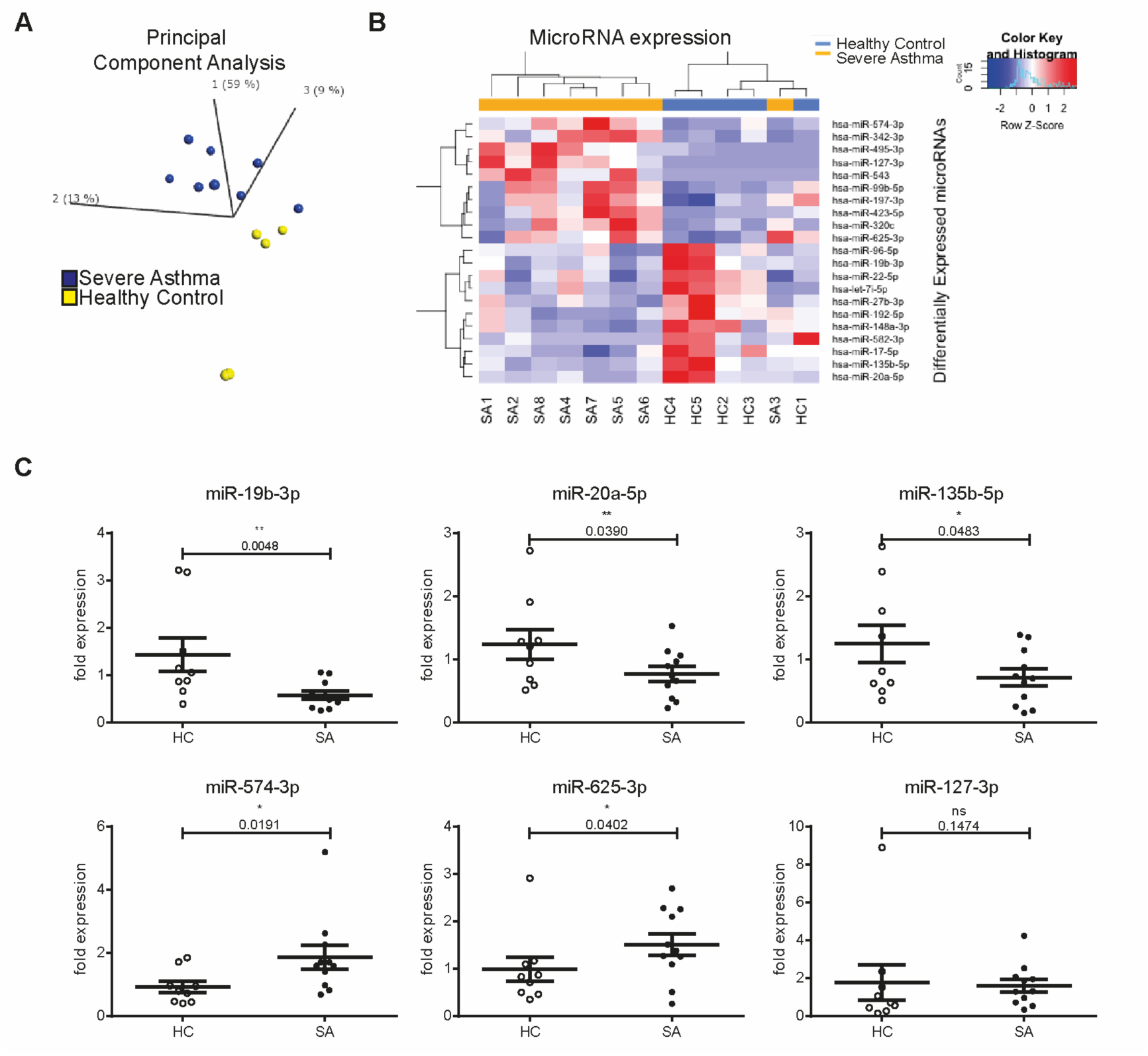
MicroRNAs are dysregulated in human severe asthma bronchial epithelium. A: Principal Component Analysis plot showing the distribution of healthy (yellow) and severe asthma (blue) samples according to the levels of differentially expressed microRNAs (P <0.05, 0.66 > SA/HC ratio > 1.5). B: Heatmap depicting unsupervised clustering of healthy controls and severe asthmatics based on the expression values of differentially expressed microRNAs (P < 0.05, 0.66 > SA/HC ratio > 1.5). C: Dot plots (mean + standard error of the mean) representing qPCR analysis of several microRNAs (n=9 HC, n= 11 SA). Statistics were done employing t-tests. HC: Healthy control; SA: Severe Asthma. * P < 0.05, ** P < 0.01.

Ingenuity Pathway Analysis (IPA) highlighted the impact of the remaining 20 dysregulated microRNAs on molecular and cellular functions of relevance to asthma (Supplementary Table S3), as well as association with *Organismal Repair and Abnormalities* (18 microRNAs), *Inflammatory Response* and *Immunological Disease* (9 microRNAs each). Thus, microRNAs dysregulated in SA may underlie important general pathological processes in asthma related to epithelial repair and inflammation via post-transcriptional mRNA regulation.

### Total mRNA expression is altered in severe asthma

As microRNAs may regulate mRNA levels by destabilization (4,24), we performed transcriptomics analysis by RNA-sequencing. Genome-wide mRNA expression analysis revealed 16,277 expressed genes in BECs (median read counts ≥ 10) with 194 Differentially Expressed Genes (DEGs, P ≤ 0.01, Supplementary Table S4). Unsupervised cluster analysis of DEGs separated HC and SA samples (Figure 2A). RT-qPCRs (Figures 2B and Supplementary Figure S1) validated a decreased expression of *COL21A1*, *CEBPA* and *CTSD* mRNAs and an up-regulation of *IGFL1*, *IL23A*, *ABCC4*, *PDPN* and *IL31RA* gene expression in SA compared to HC on an expanded cohort (n=10 HC, n=11 SA).

**Figure 2.**
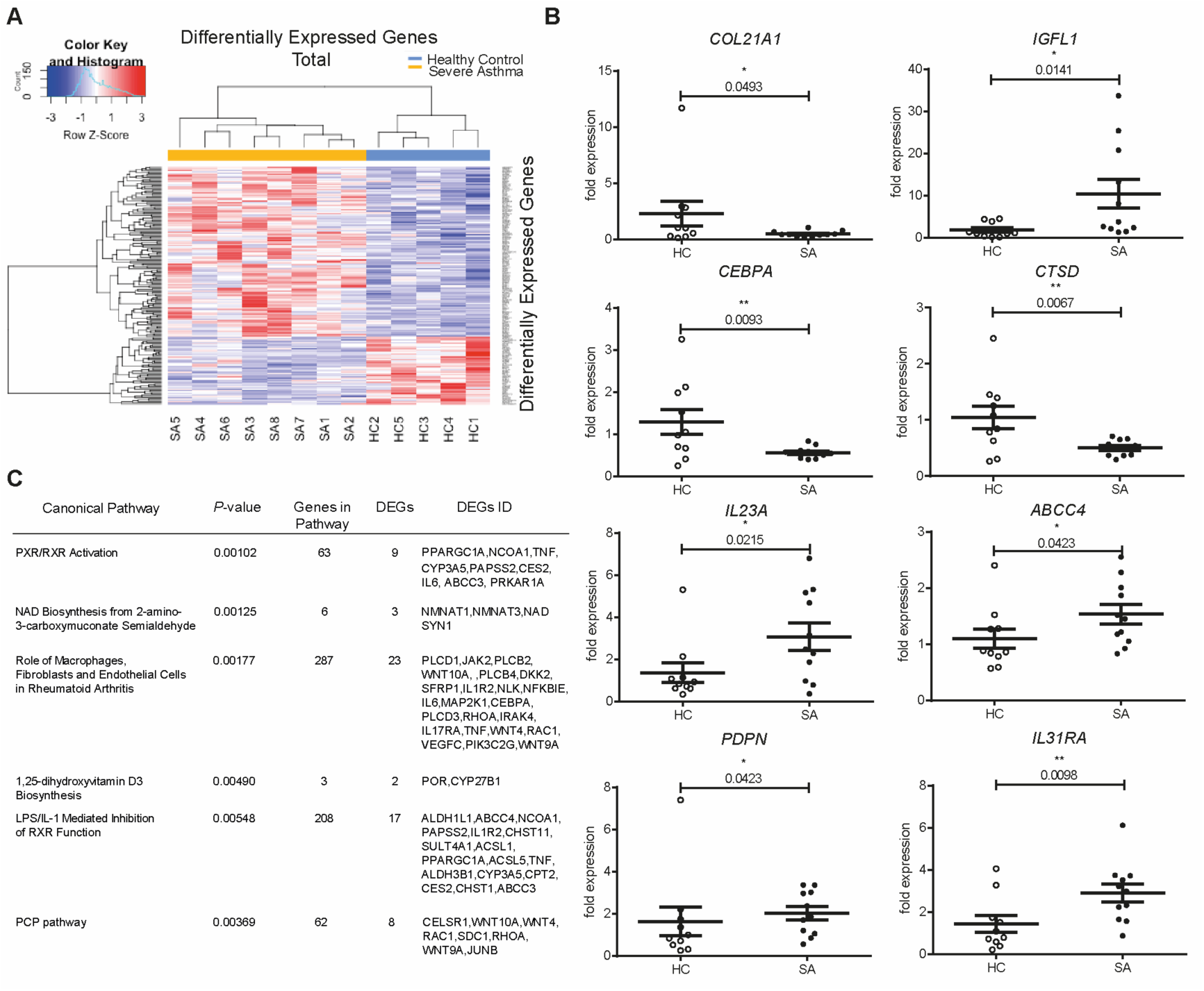
Genome-wide mRNA expression is dysregulated in severe asthma. A: Heatmap showing unsupervised clustering of Differentially Expressed Genes (P ≤ 0.01) in the Total fraction. B: Dot plots (mean + standard error of the mean) representing qPCRs validating the RNA-seq dataset (n=10 HC, n=11 SA). Statistics were done employing t-tests. C: Table showing 6 of the top 10 pathways predicted by Ingenuity Pathway Analysis for differentially expressed genes. Statistics were done employing Fisher’s 2-tailed test. HC: Healthy Control; SA: Severe Asthmatics; DEGs: Differentially Expressed Genes. * P < 0.05, ** P < 0.01.

IPA identified dysregulated pathways attributable to the detected DEGs (Figure 2C, Supplementary Table S5). This revealed non-Type2 inflammatory pathways, as well as glucocorticoid activation- and drug metabolism-related pathways. The pathway with the strongest statistically significant dysregulation (P = 0.001) was PXR/RXR (Pregnane × Receptor/Retinoid X Receptor), related to endobiotic and xenobiotic/drug metabolism (34), consistent with the high dose corticosteroid therapy prescribed to the severe asthmatics (35). Dysregulated LPS/IL-1 Mediated Inhibition of RXR Function (P = 0.0055) and 1,25-dihydroxyvitamin D3 Biosynthesis (P = 0.0049) were also evident.

### Translation in severe asthma is altered at the genome-wide level

Frac-seq was performed to determine the levels of mRNAs undergoing translation in BECs (Figure 3A) in the same healthy individuals and severe asthma patients as in Figures 1 and 2. Figure 3B depicts two representative polyribosome profiles of both healthy controls and severe asthma patients (remaining in Supplementary Figure S2). The same mRNAs as previously detected (Figure 2) were evident in the Polysome fraction (16,277 genes, median read counts ≥ 10). Severe asthmatic patients and healthy donors differed in the binding to polyribosomes of 243 genes (DBGs, *P* ≤ 0.01, Supplementary Table S6). Unsupervised hierarchical clustering of DBGs distinguished between HC and SA (Figure 3C). Noteworthy, severe asthma patients with early onset disease (onset <25 years old) clustered differently to the other five SA samples (onset >40 years old). Figures 3D and Supplementary Figure S3 show the results from the validations of several candidate genes on the expanded cohort employing qPCRs (n=10 HC, n=11 SA). This confirmed that *IGFL1*, *IL23A*, *IL1A*, *PDPN*, *IL31RA*, *ABCC4* and *LTB* genes were all more bound to polyribosomes in SA than in HC. In contrast to total mRNA, there was no evidence of decreased polyribosomal binding of *COL21A1* in severe asthma.

**Figure 3.**
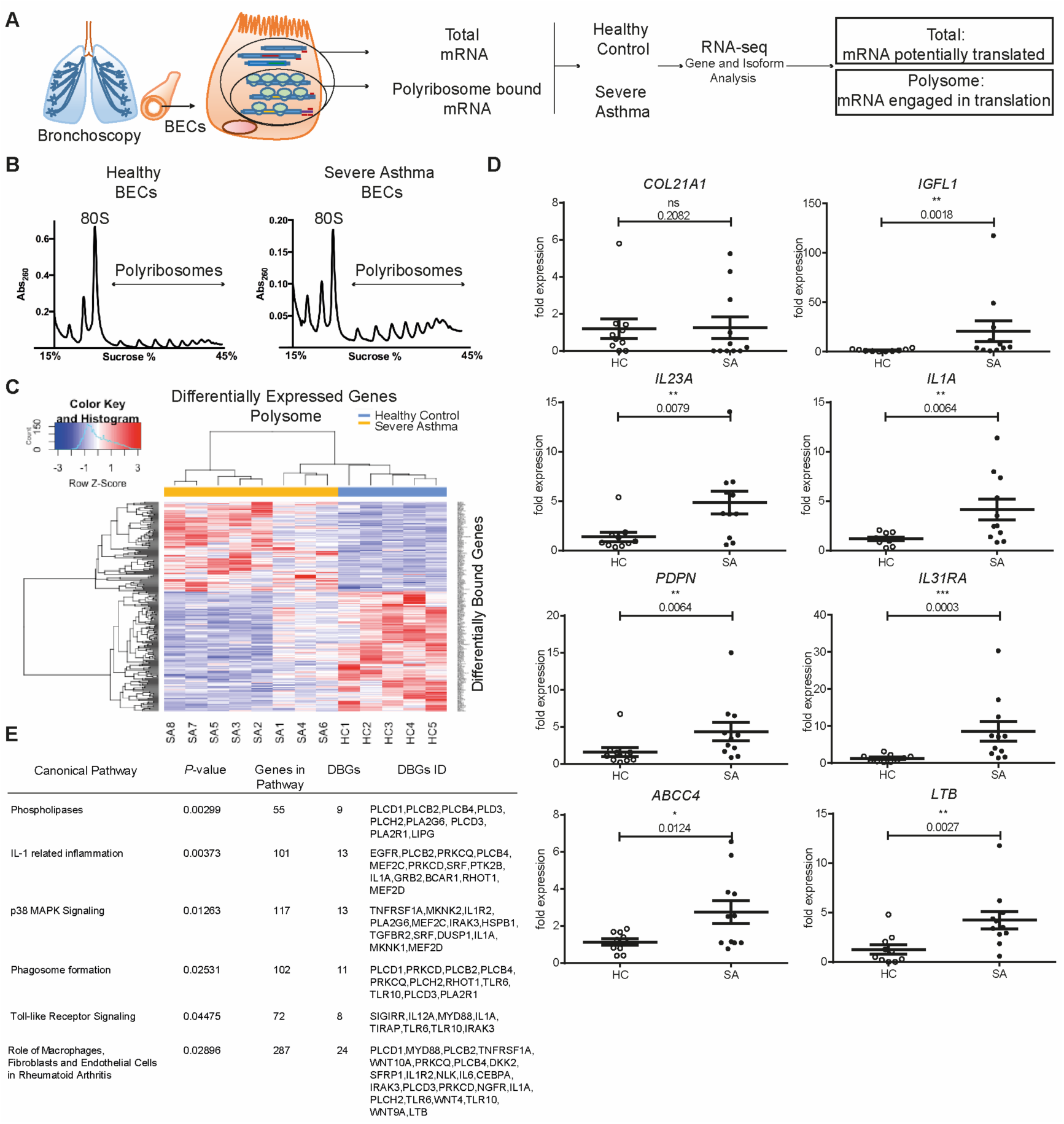
Genome-wide translation is dysregulated in severe asthma. A: Schematic of Frac-seq experiment. RNA-seq was performed on total (Total) and polyribosome-bound (Polysome) mRNA; Total HC vs Total SA and Polysome HC vs Polysome SA datasets were then compared. B. Representative polyribosome profiles from healthy (left) and severe asthma (right) BECs. C: Heatmap showing unsupervised clustering of Differentially Bound Genes (P ≤ 0.01) in the Polysome fraction. D: Dot plots (mean + standard error of the mean) representing qPCRs validating the RNA-seq dataset (n=10 HC, n=11 SA). Statistics were done employing t-tests. E: Table showing 6 pathways predicted by Ingenuity Pathway Analysis for differentially bound genes not found in the Total fraction. Statistics were done employing Fisher’s 2-tailed test. BECs: Bronchial Epithelial Cells; HC: Healthy Control; SA: Severe Asthmatics; DBGs: Differentially Bound Genes. * P < 0.05, ** P < 0.01, P < 0.001.

Pathway analysis was undertaken and dysregulated pathways clustered into the same categories as in Figure 2C (Supplementary Table S7, Figure 3E). Unlike DEGs in Total,
DBGs in Polysome did not map to glucocorticoid or endobiotic metabolism. Pathways present in Polysome DBGs and absent in Total DEGs included Toll-like receptor (TLR) signaling (*P* =0.04) and IL-1-related inflammation (*P* = 0.004), both implicated in severe asthma (36,37), as well as phagosome formation (*P* = 0.025).

Together, these results identify that there is genome-wide dysregulation of translation (mRNAs bound to polyribosomes) in asthma, and that this impacts additional pathways and genes that differ from those detected when analyzing Total mRNA levels, suggestive of underlying post-transcriptional dysregulation.

### Isoform mRNA analysis reveals structural and inflammatory anomalies in severe asthma not disclosed by aggregate gene expression

Alternative splicing analysis of the RNA-seq data revealed 30,831 mRNAs (median counts ≥ 10) in both Total and Polysome fractions. Differential Expression analysis of Isoforms (DEIs,*P* ≤ 0.01, Supplementary Table S8) revealed 319 mRNA isoforms differentially expressed between HC and SA in Total. Unsupervised clustering of DEIs clearly distinguished healthy and severe asthma samples (Figure 4A), indicating that the bronchial epithelial expression of alternatively spliced mRNAs is different genome-wide between HC and SA.

**Figure 4.**
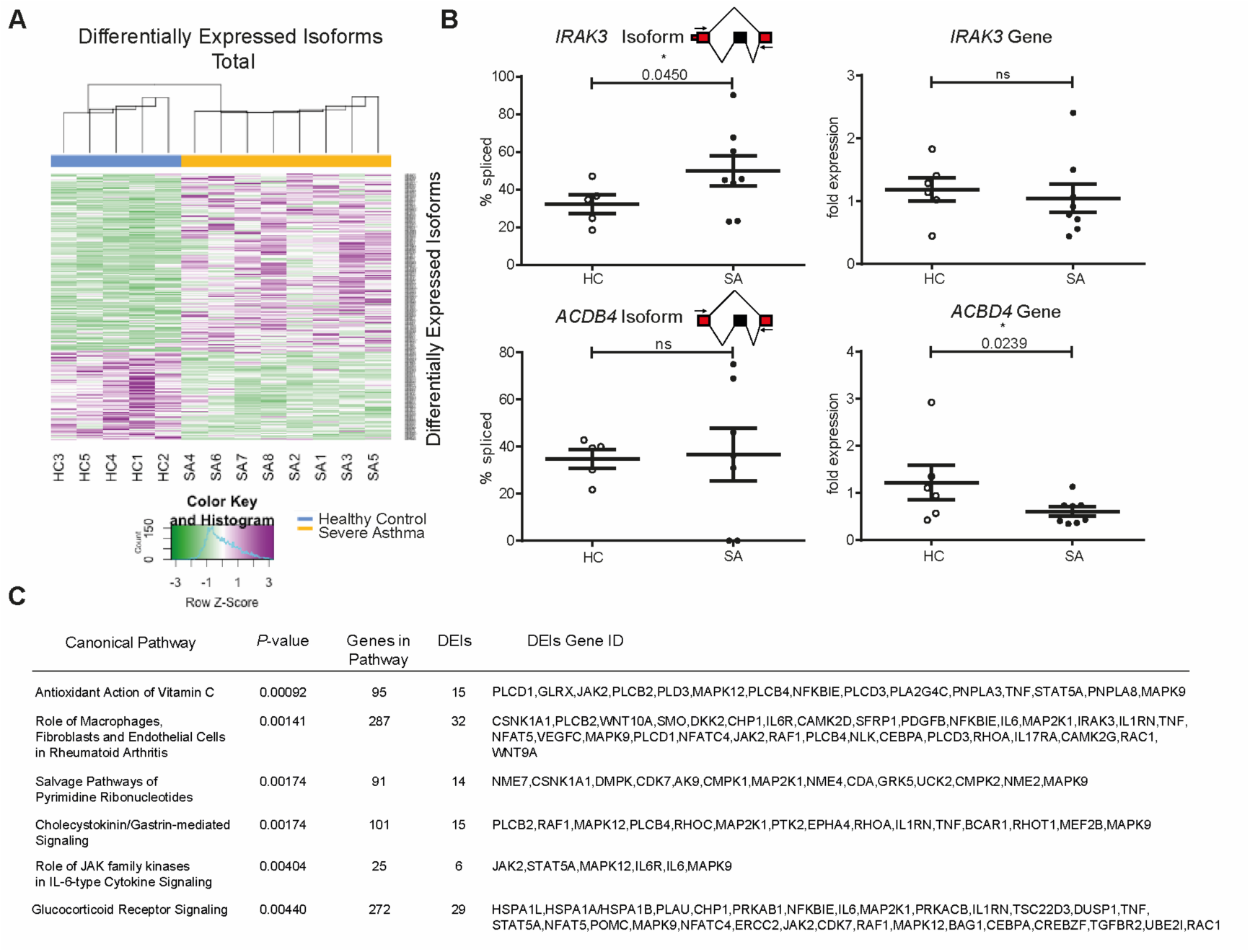
Genome-wide mRNA alternative splicing is dysregulated in severe asthma and adds novel information to that detected by gene expression analysis. A: Heatmap showing unsupervised clustering of donors according to differentially expressed isoforms (P ≤ 0.01) in the total fraction between HC and SA. B: Dot plots (mean + standard error of the mean) representing the RT-PCR splicing assays for IRAK3 and ACBD4 skipped isoforms (HC n=5, SA n=8, left column) and their corresponding qPCR assay for aggregate gene expression (HC n=6, SA n=8, right column). RT-PCRs were quantified using an Agilent Bioanalyzer DNA microfluidic chip and % splicing calculated and plotted as %inclusion= [included isoform]/([included isoform]+[excluded isoform]). Statistics were done employing t-tests. C: Table showing 6 pathways predicted by Ingenuity Pathway Analysis for differentially expressed isoforms in the Total fraction. Statistics were done employing Fisher’s 2-tailed test. DEIs: Differentially Expressed Isoforms; ns: non-significant. * P < 0.05.

The differentially expressed isoforms mapped to genes that appeared differentially expressed in Total genes (Figure 2), although alternative splicing analysis also revealed new candidates validated employing splicing assays. Figure 4B depicts some examples: left column shows the results from alternative splicing assays (RT-PCRs quantified using a Bioanalyzer) and right column those of aggregate gene expression (=all isoforms, measured by qPCRs). The skipped isoform of *IRAK3* (Interleukin-1 Receptor-Associated Kinase 3) was increased in severe asthma BECs, but was not detected differentially expressed employing aggregate gene expression assay. On the other hand, aggregate gene expression analysis detected *ACBD4* as down regulated in severe asthma while alternative splicing analysis revealed no difference. Thus, alternative splicing analysis reveals novel candidates related to asthma biology involved in inflammatory functions of bronchial epithelium not disclosed by aggregate gene expression analysis.

Differentially expressed isoforms were mapped onto pathways using IPA and grouped similarly to Figures 2 and 3 with the addition of *epithelial repair/remodeling pathways* (Supplementary Table S9) which became apparent when performing alternative splicing analysis. Figure 4C shows 6 pathways amongst the top 10 according to *P*-value. Consistently with our previous findings in Total differentially expressed genes (Figure 2C), Total differentially expressed isoforms affected glucocorticoid signalling (*P* = 0.0044), but analysis of isoforms detected new pathways including IL-6-JAK/STAT signalling (*P* = 0.004).

To interrogate the impact of alternative splicing on mRNA translation we also analyzed alternatively spliced isoforms on polyribosome-bound mRNAs. This identified 335 mRNA isoforms differentially bound to polyribosomes (DBIs) between HC and SA (Supplementary Table S10) that allowed unsupervised clustering of the samples (Figure 5A). Several candidates in the Polysome fraction were validated by splicing assays and possible differences with aggregate gene expression assessed using qPCRs (Figure 5B). The skipped isoforms of *ITGA2* and *ITGA6* genes presented increased polyribosome binding in severe asthma, information missed when performing gene expression analysis. *ITGA6* and *ITGA2* encode for integrins, key structural proteins in cell adhesion and signaling (38). *IRAK3* skipped isoform (increased only in Total DEIs, Figure 4C) showed increased binding to polyribosomes when performing analysis of aggregate gene expression.

**Figure 5.**
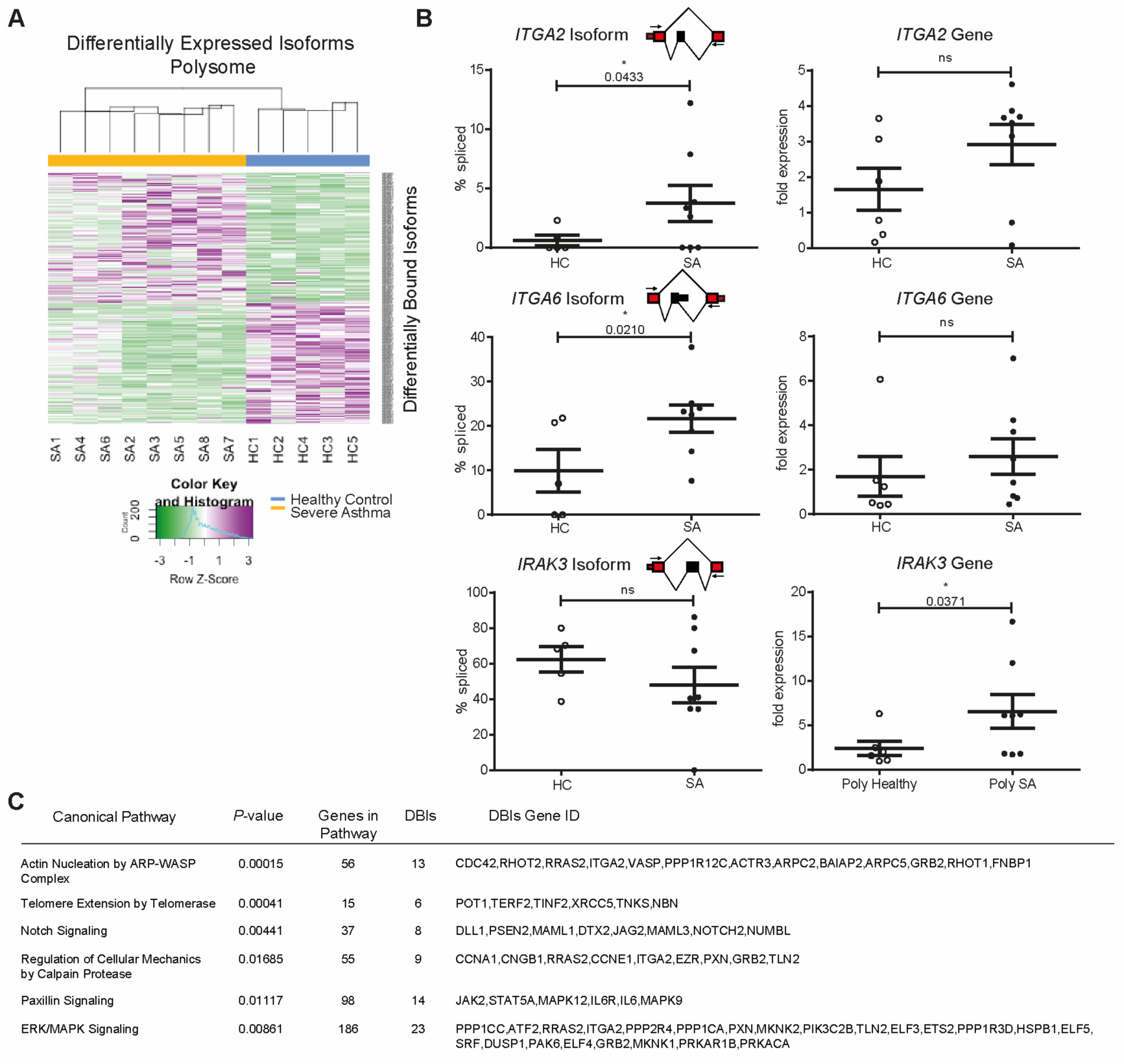
Genome-wide binding of mRNA isoforms to polyribosomes is dysregulated in severe asthma providing novel information to that detected by gene expression analysis. A: Heatmap showing unsupervised clustering of donors according to differentially expressed isoforms (P ≤ 0.01) in the total fraction between HC and SA. B: Dot plots (mean + standard error of the mean) representing the RT-PCR splicing assays for ITGA2, ITGA6 and IRAK3 skipped isoforms (HC n=5, SA n=8, left column) and their corresponding qPCR results for aggregate gene expression (HC n=6, SA n=8, right column). RT-PCRs were quantified using an Agilent Bioanalyzer DNA microfluidic chip and % splicing calculated and plotted as %inclusion= [included isoform]/([included isoform]+[excluded isoform]). Statistics were done employing t-tests. C: Table showing 6 pathways predicted by Ingenuity Pathway Analysis for differentially bound isoforms not found in the Total fraction. Statistics were done employing Fisher’s 2-tailed tests. DBIs: Differentially Bound Isoforms; ns: non-significant. * P < 0.05.

IPA of Polysome differentially bound isoforms revealed down-regulation of Notch signaling (Figure 5C and Supplementary Table S11), suggesting a decreased airway Type2-response (39) in severe asthma. Differentially bound isoforms in Polysome determined dysregulation of telomere extension, with telomere length in leukocytes previously related to asthma (40). Polysome DBIs also revealed pathways relating to epithelial cell repair/remodeling (e.g. Calpain protease or Paxillin signaling), consistent with the airways epithelium impairment observed in asthma (41).

Together, these data show that alternative splicing is dysregulated in severe asthma bronchial epithelium at a genome-wide scale affecting the translation of mRNAs encoding structural and inflammatory factors.

### MicroRNAs associate with genome-wide changes in mRNA expression at the transcriptional and translational levels

To determine the genome-wide effect of microRNAs dysregulated in the bronchial epithelium of asthma patients we aimed to identify which differentially expressed mRNAs were targeted by microRNAs on each fraction. To do so, we cross-referenced the predicted targets from TargetScan 7.1 (32) for the 20 microRNAs (Figure 1, all except for miR-127-3p) with the differentially expressed isoforms in Total and Polysome (Figures 4 and 5), as microRNAs are known to target specific isoforms (42). This showed that microRNAs significantly modify the levels of both cytoplasmic and translating mRNAs (Supplementary Figure S4). The overlap between targets in the cytoplasmic and polyribosome bound mRNA fractions was very low (Figure 6A, 11.2 % of Total DEIs and 7.6 % of Polysome DBIs), suggesting that microRNAs dysregulated in asthma may have different effects depending on the mechanism of action (mRNA degradation or blocking of translation). MicroRNAs showed preferential targeting of polyribosome-bound mRNAs (172/288) compared to cytoplasmic mRNAs (116/231) (Supplementary Figure S5A, 2-tailed Fisher’s exact test *P* = 0.0331) consistent with mRNAs presenting more or less MREs according to their association with polyribosomes (Supplementary Figure S5B and (20). These results suggest that dysregulated microRNAs in asthma may have more impact in protein translation than on mRNA levels.

**Figure 6.**
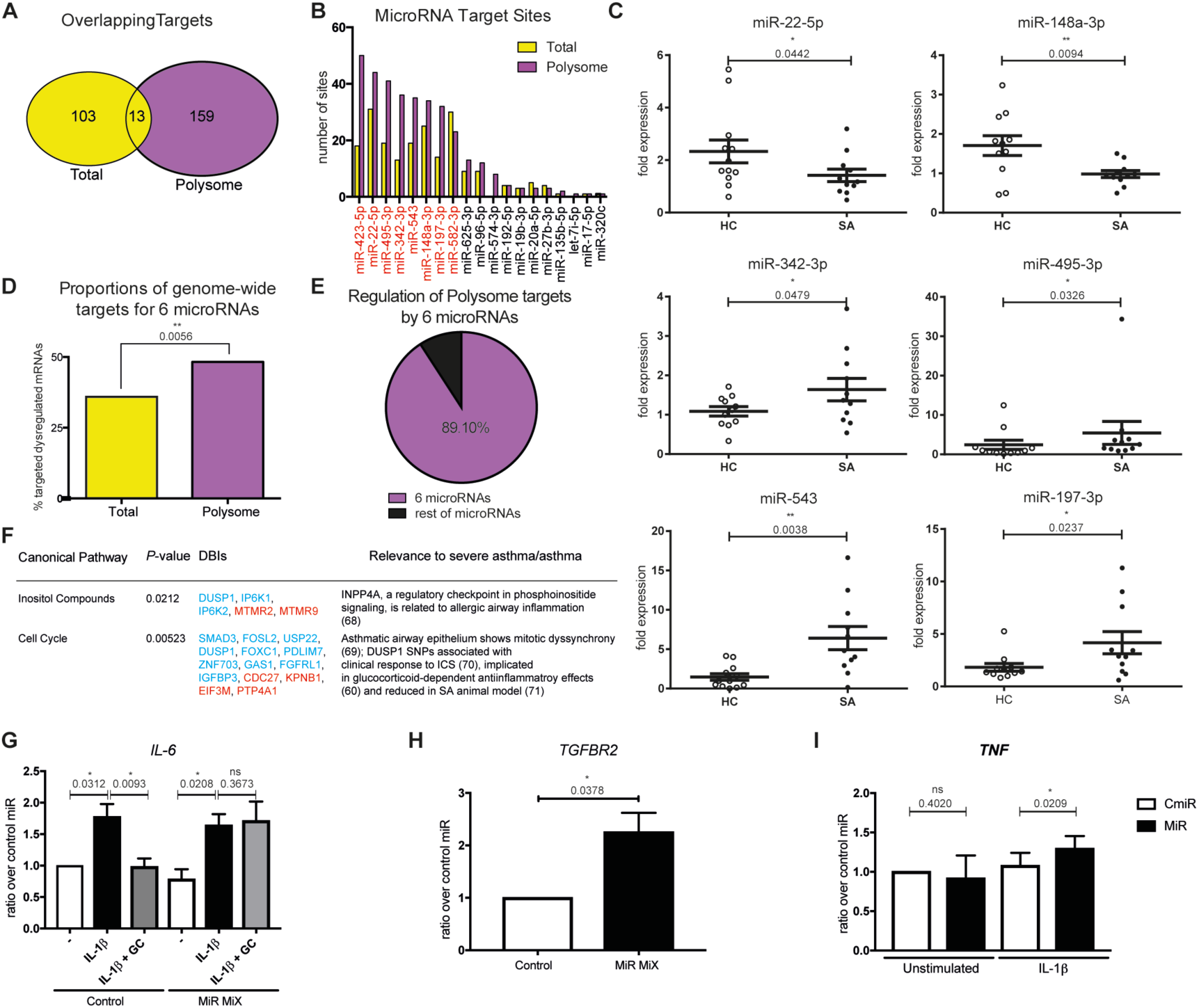
A subset of 6 microRNAs controls most of the mRNA targeting in severe asthma bronchial epithelium. A: Venn diagram depicting the overlap between total and polyribosome-bound microRNA targets by cross-referencing differentially expressed/bound isoforms with differentially expressed microRNAs in severe asthma. B: Bar plot depicting the number of targets for each differentially expressed microRNA in the total (yellow) and polyribosome bound (pink) fractions. Highlighted in red are microRNAs with most abundant sites. C: Dot plots (mean + standard error of the mean) representing qPCRs validating the subset of microRNAs amongst those with the highest number of targets in both fractions (n=11 HC, n=11 SA). Statistics were done employing t-tests. D: Bar plot depicting the proportional abundance of targets for the 6 validated microRNAs in total (yellow) and polyribosome bound (pink) fractions. Polyribosome bound mRNAs have a higher abundance of microRNA targets (2-tailed Fisher’s exact test). E: Pie chart depicting the proportion of targets amongst mRNAs controlled by microRNAs in the polyribosome bound fraction that is potentially regulated by the validated 6 microRNAs. F: Table showing the main asthma-related pathways predicted by Ingenuity Pathway Analysis for mRNA isoforms differentially bound to polyribosomes and potentially targeted by the hub of 6 microRNAs. Red: up-regulated; blue: down-regulated. Statistics were done employing Fisher’s 2-tailed test. G, H, I: Bar plots showing the results from transfecting the hub of microRNAs onto healthy BECs. 48h post-transfection cells were pre-treated or not during 2h with dexamethasone, stimulated or not with IL-1β and harvested 24h later. * P < 0.05, ** P < 0.01.

The relevance of each one of the 20 microRNAs in targeting Total and Polysome mRNAs was then evaluated. Figure 6B shows the number of targets per microRNA, which revealed that a group of only 8 microRNAs controls most of the mRNA changes detected in severe asthma both at cytoplasmic and polyribosome binding levels. Amongst those 8 microRNAs, dysregulated expression of 6 microRNAs was validated in our larger cohort of patients (n=11 for each group, Figures 6C and Supplementary Figure S6). These 6 microRNAs were also predicted to preferentially modulate changes on translating mRNAs (48.2% as opposed to 35.9% in Total, Figure 6D, *P* = 0.0056). Network analyses of microRNA :: target interactions showed that the target mRNAs of these 6 microRNAs are in many cases co-regulated by multiple microRNAs. This suggests that the expression of these mRNAs is tightly controlled, but disrupted in severe asthma patients (interactive networks in Supplementary Interactive Figures S1 and S2). Strikingly, this network of 6 microRNAs was predicted to control almost 90% of all targeted mRNAs in the Polysome fraction (Figure 6E), which mapped to inositol pathways and cell cycle (Figure 6F, Supplementary Table S12) the latter desynchronized in asthma (43).

In order to validate our findings, we transfected healthy bronchial epithelial cells with the microRNA network to evaluate the effects of the dysregulated microRNA hub found in severe asthma BECs. Namely, anti-miR-22-5p, anti-miR-148a-3p, pre-miR-342-3p, pre-miR-495-3p, pre-miR-543 and pre-miR-197-3p oligonucleotides were co-transfected at equimolar concentrations and compared to BECs co-transfected with negative anti-miR and pre-miR controls at the same concentrations. Cells were then treated with IL-1 β, which is found up-regulated in SA (and highlighted in our pathway analysis) as well as assessed for glucocorticoid sensitivity. Figures 6G-I show that the microRNA network was able to ablate the inhibition by glucocorticoids of IL-1 β driven IL-6 mRNA expression (Figure 6I). The microRNA hub was also able to up-regulate *TGFBR2* mRNA expression (Figure 6H) as well as significantly increase IL-1β-driven *TNF* mRNA expression. These are all characteristics well defined in severe asthma patients, highlighting the biological importance of microRNA dysregulation in severe asthma bronchial epithelium.

Integrated together, our data demonstrate that there is genome-wide deregulation of post-transcriptional processes in human asthma. Our work shows that the dysregulation of a microRNA hub causes genome-wide dysfunction in the translation of mRNAs encoding structural and inflammatory factors in the bronchial epithelium of severe asthma patients, and demonstrates the value of integrating multiple–omics datasets when investigating human biological processes.

## DISCUSSION

Our work integrating Frac-seq and small RNA-seq reveals that microRNAs, cytoplasmic mRNAs and translating mRNAs are all dysregulated in human primary airway cells from asthma patients. Integrating differentially expressed microRNAs and mRNAs determines that altered microRNA expression impacts on the detected mRNA changes, underlying abnormalities relating to inflammation, glucocorticoid sensitivity and epithelial repair detected by pathway analysis. This is effected through genome-wide modulation of mRNA levels and most predominantly through regulation of mRNA binding to polyribosomes. To our knowledge, ours is the first study to employ polyribosome profiling in human clinical samples and demonstrates that this approach reveals disease pathways and mRNA candidates not disclosed by other approaches, mainly transcriptomics, widely employed when studying human disease. For example, severe asthma patients with an early disease onset clustered differently than those with later onset (Figure 3) in the Polysome fraction. Although more numbers are needed to confirm this observation, it is consistent with reports suggesting that early and late onset SA represent stratified types of asthma with different etiologies (44). Polyribosome-bound mRNA analysis therefore highlights disease-related information overseen by Total mRNA measurements and may serve as a novel tool to endotype patients, key into understanding and managing patients with complex diseases such as asthma and one of the hallmarks of personalized medicine.

One of the key advantages of Frac-seq is that it has more coverage than current proteomics approaches and informs about underlying molecular mechanisms of mRNA regulation, revealing mRNA isoforms preferentially bound to polyribosomes (20). Additionally, it reveals alternative splicing events that may lead to changes in 3’UTRs or 5’UTRs, which may not render differences in the aminoacid sequence of proteins but strongly impact on mRNA translation regulation. Expression changes in mRNAs undergoing translation reveal pathway abnormalities in severe asthma relating to Toll-like receptor signaling, the IL-1 pathway and to p38 MAPK. These are all pathways distinct from those related to classical type 2 inflammation described in untreated steroid-responsive asthma (45) and mostly absent in our cohort (Supplementary Figure S7). Moreover, the dysregulated microRNA network was found to regulate IL-1 β responses in BECs (Figure 6G and 6I), highlighting the intricate post-transcriptional dysregulation underlying disease characteristics in severe asthma BECs.it is estimated that all human genes undergo alternative splicing (AS) producing at least two alternative mRNA isoforms (19), with around 80% of AS events estimated to lead to protein modifications (46) and AS influencing protein output (21). Moreover, 25% disease-related mutations have been linked to defects in splicing (47). If such changes existed in asthma, they could have profound and genome-wide implications in the proteins expressed by cells and thus in cellular function (21). The more in depth analysis of mRNA isoforms differentially bound to polyribosomes identified, amongst other findings, defective signaling related to epithelial repair/remodeling pathways (e.g. Calpain protease or Paxillin signaling), supporting evidence from other approaches that there is an altered epithelial repair phenotype in severe asthma (10,48). These results are also supported by increased binding to polyribosomes of *ITGA2* and *ITGA6* mRNA isoforms in severe asthma (Figure 5), integrins being key adhesion and signaling proteins in the barrier (38). Moreover, our data supports that selective binding of mRNA isoforms to polyribosomes is, at least partially, determined by microRNA targeting (Supplementary Figure S5B).

The relevance and implications of employing Frac-seq in human clinical samples are also supported by previous findings at the protein level. The identification of increased epithelial polyribosomal *EGFR* mRNA binding in severe asthma (Supplementary Figure S8) supports the reported increased epithelial expression of EGFR protein (49). This has been attributed to a repair phenotype promoted by TGFβ-induced cell cycle inhibition (50). The present finding of increased binding to polyribosomes of *TGFBR2* (Supplementary Figure S8), the main epithelial receptor for TGFβ, regulated by the microRNA hub (Figure 6H), would underlie this potential. Bacterial products induce inflammatory responses via EGFR and TLR-mediated pathways (51), with TLR-signaling present in pathways altered in Polysome, and viruses and bacteria exploiting EGFR to facilitate their survival (52,53) or to attenuate the host response (54). Increased *IL23A*, *IL31R* and *IL1A* mRNA translation in the epithelium would be consistent with an immune system orientated towards Type I and type 17 directed inflammation (Supplementary Figure S7), more characteristic of a bacterial driven process, as would be the Toll receptor pathway activation, phagosome formation and the evidence of altered *IL6* and *IL6R* isoforms binding to polyribosomes (Supplementary Table S10).

The translation of genes linked to innate immune responses raises the possibility that they arise in response to the altered airway microbiome in severe asthma (55). An altered airway microbiome may also contribute to steroid resistance, one of the features of severe asthma. Patients with severe asthma suffer from persistent disease despite use of maximum therapy, including high dose of glucocorticoids. In the present study, glucocorticoid signaling was detected in Total, suggesting that corticosteroids are effectively reaching the airways of these patients and indicating that the lack of effect of steroid treatment is not a reflection of lack of adherence to treatment. Consideration needs to be given as to whether some of the differential expression changes in severe asthma reflect steroid treatment. This does not appear the case for microRNAs as previous *in vivo* studies have reported that glucocorticoids have little effect on microRNA expression in bronchial epithelium (56,57). Whilst steroids affect mRNA expression, there was no overlap between the identified up-regulated differentially expressed genes in Total or Polysome and gene changes described from previous genome-wide studies aimed at determining the effect of corticoids on gene expression (58,59). Thus the majority of described differential gene changes are likely to reflect alterations related to the underlying disease pathophysiology rather than directly due to their current therapy. Whilst glucocorticoid signaling was evident in Total, no glucocorticoid signaling was detectable in the polyribosome-associated pathways. *DUSP1* was identified as down-regulated in SA, whilst several studies have highlighted the importance of this in glucocorticoid anti-inflammatory effects (60,61). Indeed, we found that the microRNA network ablated the inhibition by glucocorticoids of IL-1 1β-driven IL-6 mRNA expression (Figure 6G), potentially mimicking SA insensitivity to corticosteroids. Thus in severe asthma there are nonsteroid responsive pathways evident within the bronchial epithelium mRNA translational signature, and regulated by the hub of 6 microRNAs- a novel finding of potential clinical relevance solely disclosed by our approach. Together these data highlighting the need for the development of additional or alternative approaches to therapy in these patients and the importance of using of polyribosome profiling to reveal novel pathophysiological mechanisms, of potential implications in other inflammatory diseases as previously implied in cancer (62). Integrating small RNA expression allowed us to associate changes between mRNA and microRNA levels. Whilst previous studies have demonstrated the relevance of individual microRNAs in disease and particularly in asthma (63-65), there are no genome-wide studies addressing the relationship of microRNA levels with cytoplasmic and translational mRNA changes in asthma, and to our knowledge, in any other disease employing human clinical samples. Our results show that microRNA effects mostly rely on a hub of six microRNAs that potentially regulates ∼ 50% of all dysregulated mRNAs undergoing translation and ∼35% of cytoplasmic mRNA changes detected in asthma patients. We considered 6-7-and 8-mer MREs, consistent with previous studies showing that all these contribute to target abundance (66). Our results are also consequent with microRNAs affecting both mRNA degradation (4,24) and translation inhibition (2,5). Recent work showing the relevance of ribosome binding for microRNA action (5), as well as mRNA degradation happening co-translationally (67) support our observation that microRNAs preferentially modulate mRNAs bound to polyribosomes. We could not find a relationship between the number of mRNA targets and the expression levels of the dysregulated microRNAs, as previously noted by (62) in neuroblastoma. MiR-148a-3p, with the highest expression in healthy, and miR-197-3p, with the highest expression in asthma, had the fewest number of mRNA targets amongst the hub of six microRNAs. Our results add novelty on a broad spectrum of biological contexts, as we have considered physiological changes in microRNA and mRNA levels in disease, rather than studying the isolated behavior of individual microRNAs or target mRNAs. In this context, microRNAs seem to target different mRNA populations when analyzing cytoplasmic or polyribosome-bound mRNAs (Figure 6). We cannot exclude the possibility that inhibition of translation may have precluded cytoplasmic changes due to decay (4), or indeed, that some of the up-regulation observed for microRNA targets may be due not only to a release of inhibition from microRNAs down-regulated but also to transcriptional activation. Our results add to the latest studies demonstrating the intricate relationship between ribosome binding and mRNA fate with regards to stability and microRNA action.

In conclusion, our work demonstrates that whilst translation and alternative splicing are well-controlled processes in health they are genome-wide dysregulated in asthma patients, and that microRNAs account for many of the observed mRNA changes. This study reveals a new role for microRNAs in controlling impaired translation in asthma of potential implications in other diseases, placing a hub of six microRNAs as potential future therapeutic candidates to address steroid unresponsive epithelial activation in asthma. Our approach demonstrates the feasibility and essential importance of studying post-transcriptional gene regulation when investigating human biology and disease.

## MATERIALS AND METHODS

Full details are in Supplementary Materials and Methods.

### Study volunteers and consent to participate

Non-smoking volunteers aged 18-65 years were recruited from the Wessex Severe Asthma Cohort and age/sex matched healthy controls from a departmental database (Supplementary Table S1). All participants gave written informed consent.

### Bronchial epithelial brushings and cell culture

Flexible bronchoscopy was undertaken as previously described (25) and brushings obtained from the right *bronchus intermedius* using disposable, sheathed bronchial brushes (Olympus BC-202D-1210). Brushings were spun at 1200 rpm 10 minutes, medium discarded and cells resuspended in complete BEGM (Lonza, Blackley, United Kingdom). Bronchial epithelial cells were cultured as previously described (7). Briefly, bronchial epithelial cells were cultured in collagen (Thermo Fisher Scientific, Loughborough, UK)-coated T25 flasks in BEGM complete medium (Lonza, UK) and passaged onto 15cm^2^ dishes when 80% confluent. All sequencing experiments were done in passage 1 cells; microRNA transfections were done in p3/p4 bronchial epithelial cells.

### Small RNA-sequencing

Libraries and small RNA sequencing were done in Ocean Ridge (Florida, USA). Libraries were made employing NEBNext Small RNA-Seq Library Preparation Kit (New England Biolabs-Ipswich, USA) according to manufacturer’s instructions and purified using a gel-based extraction. Libraries were sequenced with 50-bp single-end reads plus index read (TruSeq Rapid SBS Kit-HS 200 Cycle, Illumina Inc., San Diego, CA). An average of 4.6M reads passing trimming and minimum count filter (≥ 5 reads) was generated per sample. Nonredundant sequences were then aligned to genomic (hg19) and mRNA sequence (hg19) using bowtie2 (26); sequences with perfect match and 1nt mismatch were retained for further analysis. The genome-mapped sequences were further aligned to precursor and mature microRNAs in miRBase 21.0 (27) using OMAP# alignment software developed at Ocean Ridge Biosciences. To facilitate statistical analysis, the raw reads were converted to reads per million (genome) mapped reads (RPM). Mature microRNA RPM values were normalized using the following formula: raw count / perfectly matched microRNA reads X 1,000,000. Tables were filtered to retain a list of annotated RNAs having a minimum of 10 mapped reads (Detection Threshold) in 25% of samples; for these filtered read tables, missing values were replaced with the average RPM value equivalent to 1 read. Library packages are available on CRAN. MicroRNA clustering in Figure 1 was done in R using kendall/ward.D method. Further details are in Supplementary Materials and Methods.

### Polyribosome profiling

Polyribosome profiling was done as described previously (20,22) with the addition of 500μg/mL cycloheximide (Sigma-Aldrich, Dorset, UK) in the lysis buffer, in passage 1 BECs. The 80S-to-polyribosome ratio is a signature of the translational status of cells and it is greatly modified when global translation is increased or impaired (28). We did not observe a difference in the translational profile between HCs and SA, as shown by similar ratios of monosome (80S) to polyribosome peaks (Figure 3B and Supplementary Figure S2).Polysome excludes the monosomal fraction (80S) as this may contain mRNAs not undergoing translation or lowly expressed genes (28,29). RNA was isolated from the individual polyribosomal peaks, pooled to constitute the Polysome fraction and sequenced. Details of the protocol and buffer composition are in Supplementary Materials and Methods.

### RNA-sequencing

RNA quality was assessed using a Bioanalyzer (Agilent) with all sequenced samples showing a RIN value greater than 7 (RIN average +/-standard deviation =8.95 +/-0.62). Libraries and sequencing were done in Expression Analysis Inc. (Durham, USA). Libraries were made using TruSeq Stranded mRNA Library Prep Kit (Illumina Inc., San Diego, CA) and sequenced in a Hiseq 2500 (Illumina Inc., San Diego, CA) platform (100bp, paired-end sequencing). A minimum of 14M reads/sample was generated. RSEM v1.2.0 was used to quantify and compute estimated counts by genes and transcripts (30) using the UCSC knownGene. All 77k isoforms defined by UCSC hg19 were considered as initial candidates. 12,485 isoforms are associated with only one gene, and thus were not considered further. Over 12k isoforms had 0 or nearly 0 counts for all subjects. Upper Quartile normalization was used to normalize between different samples with different read depths for both aggregate gene expression and isoform analysis. Each sample was scaled to that the Upper Quartile of counts was equal to 1000. Only genes and isoforms with median counts of 10 or more on each group (Total or Polysome) were taken into account. PROC-GLM was used to perform preliminary statistical analysis. Clustering methods employed were: in heatmaps for differentially expressed genes (Figures 2 and 3) Pearson/mcquitty method and in heatmaps for differentially expressed isoforms (Figures 4 and 5) using Kendall/centroid method. Heatmaps were done in R. Further details are in Supplementary Materials and Methods.

### RT-qPCR and Splicing Assays

MicroRNA validations were performed using the miScript system (QIAGEN, Manchester, UK) following manufacturer's instructions. MicroRNA expression was normalised against hsa-let-7a-5p.

RNA was retro-transcribed using High Capacity cDNA Reverse Transcription Kit (Thermo Fisher Scientific, Loughborough, UK). Validations for differentially expressed genes and microRNA effects (Figures 2, 3, 4, 5 and 6) were performed employing qPCRs using TaqMan Gene Expression Assays (Thermo Fisher Scientific, Loughborough, UK), using the primers with maximum coverage. Supplementary Figure S8 *EGFR* primers were kindly provided by N. Smithers and designed by Dr D. Smart (both at University of Southampton) and used as a SYBR Green assay (GoTaq qPCR Master Mix from Promega, UK). Details of primer sequences are in Supplementary Materials and Methods. Gene expression was normalised against *GAPDH.* Splicing was measured employing a Bioanalyzer 2100 (Agilent) and % spliced calculated as Included/(Included+Skipped).

### Pathway Analysis

Data were analyzed using QIAGEN’s Ingenuity Pathway Analysis (IPA, QIAGEN, www.qiagen.com/ingenuity), employing genes/isoforms with a cut-off of *P* ≤ 0.05 (Benjamini-Hochberg *P*-adjusted ≤ 0.05, (31). Analysis with a cut-off *P* ≤ 0.01 missed important disease-related information. Grouping of pathways was done employing the IPA database and bibliography.

### MicroRNA Analysis

MicroRNA :: target interaction in Figure 6 was determined by cross-referencing the up regulated differentially expressed isoforms (as gene ID) with the down regulated microRNAs and *vice versa* employing TargetScan 7.1 (32). We increased the stringency by accounting only for isoform ratios (SA/HC) of more than 1.5 or less than 0.66. Importantly, when sliding the ratios to less than 0.5 or more than 2 as well as not adding any restrictions on ratios, our results relating to the mRNA expression cumulative distributions differences remained significant (Supplementary Figure S4). Interactive networks in Supplementary Interactive Figures S1 and S2 were done employing networkD3 package in R.

### Image processing

Graphs for polyribosome profiles, validations, pathway analyses and microRNA analyses were produced using GraphPad Prism version 6.07 or v7, GraphPad Software, San Diego, CA, USA, www.graphpad.com. Heatmaps were produced using R statistical language, version 3.2.0 (R project for Statistical Computing, https://www.r-project.org/). All packages used in R are available in CRAN repository. All figures panels were put together using Adobe Illustrator CS6. Frequency distribution plots in Supplementary Figure S4 were done by adjusting the data to a non-linear regression to facilitate and clarify data presentation employing GraphPad Prism v6.07.

### Statistical analysis

Clinical parameters statistics (except for sex and atopy) were done employing a MannWhitney U-test for non-parametric data and unpaired t-tests for parametric data comparisons, according to D’Agostino & Pearson omnibus normality test. Differences in sex and atopy between HC and SA were tested employing a Fisher's exact test. Statistical analysis was performed using GraphPad Prism version 6.07, GraphPad Software, San Diego, CA, USA, MicroRNAs shown inwww.graphpad.com.

MicroRNAs shown in Figure 1A and 1B had a *P* ≤ 0.05 and a restricted fold change of less than 0.66 or more than 1.5 (SA/HC), which showed an FDR ≤ 0.05 when corrected using the Benjamini-Hochberg method (31). For gene and isoform expression, a restrictive *P* ≤ 0.01 (two group t-test) was taken to perform the heatmaps in Figures 2-5. Nominal P-values from RNA-seq datasets (two-group test, *P* ≤ 0.01 and *P* ≤ 0.05, Supplementary tables S4-S11) were corrected by applying the Benjamini-Hochberg method (31) employing R, which showed an FDR ≤ 0.05 for all genes and isoforms displayed in Figures 2-5 and used in pathway analysis. FDR was calculated also experimentally based on the gene expression assays (0.136), and was in the 0.1-0.2 range in accordance with previous studies (20,33). The *P*-value displayed in all pathway analyses was calculated in IPA employing a Fisher’s Exact test. All statistical analyses for validations were performed using GraphPad Prism employing a Mann-Whitney U-test for non-parametric data and unpaired t-tests for parametric data comparisons, according to D’Agostino & Pearson omnibus normality test. Statistical analysis of proportions for microRNA targeting between Total and Polysome (Figure 6) was done employing the Fisher's exact test in GraphPad Prism version 6.07. Statistics comparing cumulative distributions in Supplementary Figure S4 were performed employing a Kolmogorov-Smirnov test as in (32) using GraphPad Prism version 6.07, GraphPad Software, San Diego, CA, USA, www.graphpad.com.

All packages used in R are available in CRAN repository.

Interactive MicroRNA :: target networks are supplied as Supplementary Interactive Figures S1 and S2.

## ACKNOWLEDGEMENTS

The authors are grateful to the asthmatic and healthy participants who voluntarily participated in the research and thank the nursing support from the Southampton NIHR Respiratory Biomedical Research Unit and Southampton Centre for Biomedical Research (SCBR) that enabled the study. The authors thank Prof Mariana Castells, Dr Christopher Woelk, Dr Yawwani P Gunawardana, Dr Jeremy R Sanford, Dr Aishwarya Griselda Jacob and Dr Michael Breen for critical input to the manuscript. The authors thank Dr Michael Breen for statistical and computational analysis advice and Dr Jeongmin Woo for advice in R. The authors thank Dr Michael Edwards for his help with the experiments to test microRNA effects.

This work was supported by Medical Research Council UK [grant numbers G0800649 Wessex Severe Asthma Cohort, MR/K001035/1, G0900453]. Equipment (PHH and TSE) and reagents (RTMN) grants from Southampton Asthma, Allergy and Inflammation Research (AAIR) Charity. RTMN was in receipt of Post-doctoral Career track award from the Faculty of Medicine, University of Southampton.

## AUTHOR CONTRIBUTION

RTMN conceived, performed and analyzed the experiments; HR performed bronchoscopies, supervised clinical input; M.P. and R.C.C. contributed to the experimental work; MN contributed to the computational and statistical analysis; TSE and PHH conceived and analyzed experiments. All authors contributed to scientific discussions and manuscript writing and approved the final draft.

## THE PAPER EXPLAINED

### PROBLEM

Severe asthma is a chronic inflammatory disease of the airways characterised by poor response to therapy. The bronchial epithelium is key in asthma, but the underlying pathophysiological mechanisms remain poorly understood. Post-transcriptional gene regulation is poorly studied in asthma and may give novel insight into disease mechanisms.

### RESULTS

We performed paired microRNA, cytoplasmic and polyribosome-bound (Frac-seq) RNA-sequencing in bronchoepithelium isolated from clinical samples from healthy donors and severe asthmatic patients. Our data demonstrate that post-transcriptional gene expression is genome-wide dysregulated in bronchial epithelium from severe asthmatics. Severe asthmatics present genome-wide microRNA and alternative splicing dysregulation, with alteration of specific mRNA isoforms binding to polyribosomes that differs from those dysregulated at the cytoplasmic level. We reveal a hub of six microRNAs potentially controlling ∼90% of all microRNA targeting on these cells, showing preferential targeting of mRNAs undergoing translation. Alteration of this network in healthy bronchoepithelium mimics severe asthma characteristics.

### IMPACT

Our data demonstrate that mRNA splicing and translation are genome-wide dysregulated in severe asthma. Polyribosome-bound mRNA analysis reveals novel pathophysiological pathways in airways epithelium not disclosed by the more commonly used transcriptomics approaches. Integrating our data we demonstrate that a small hub of six dysregulated microRNAs renders bronchial epithelial cells insensitive to corticosteroids, one of the hallmarks of severe asthma. Our novel approach highlights the essential importance of assessing post-transcriptional gene regulation when investigating asthma and potentially other inflammatory-related diseases.

## DATA AVAILABILITY

The sequencing datasets are stored in the NCBI Gene Expression Omnibus repository (http://www.ncbi.nlm.nih.gov/geo/) as GSE85216, GSE85215 and GSE85214. Reviewer links are available.

